## Supplementary Table S1 for "Examining heterogeneity in dementia using data-driven unsupervised clustering of cognitive profiles"

| **Test Name** | **Interpretation/meaning** | **Range** |
| --- | --- | --- |
| Boston Naming Test | Assesses an individual’s ability to name pictured objects and measures language ability, specifically confrontational word retrieval. | Naming difficulty  >45 : None  35-45: Mild  <35 : Significant |
| Mini-Mental State Exam | A brief 30-point questionnaire that is used to screen for cognitive impairment and to estimate the severity and progression of cognitive decline. | Impairment level  24-30 : None  18-23 : Mild  10-17 : Moderate  0-9 : Severe |
| Short Blessed Test | A brief assessment tool for detecting cognitive impairment and dementia, focusing on memory, orientation, and concentration. | Impairment level  0-4 : None  4-10: Questionable  >10 : Significant |
| Word List Memory Task | Evaluates verbal memory by having the individual recall a list of words over several trials and after a delay. | 0 - 30  (higher = better) |
| Verbal Fluency | Assesses executive function and language abilities by requiring the individual to generate as many words as possible from a given category or starting with a specific letter. | 0 - 15  (higher = better) |
| CDR - Memory | Measures the degree of memory loss and its impact on daily functioning. | Impairment level  0 : None  0.5: Questionable  1 : mild  2: moderate  3: severe |
| CDR - Orientation | Assesses an individual's awareness of time, place, and personal context. |  |
| CDR - Judgment and Problem Solving | Evaluates the ability to handle complex and abstract tasks, solve problems, and make sound decisions. |  |
| CDR - Community Affairs | Measures the involvement and independence in community activities and responsibilities. |  |
| CDR - Home and Hobbies | Assesses engagement and performance in household tasks and personal hobbies. |  |
| CDR - Personal Care | Evaluates the ability to maintain personal hygiene and carry out activities of daily living independently. |  |
