## Supplementary Table S2 for "Examining heterogeneity in dementia using data-driven unsupervised clustering of cognitive profiles"

| **Brain-related disorders** | **ICD 10 codes** | **ICD 9 codes** |
| --- | --- | --- |
| Memory Loss | R41.3, R41.89, G31.84, R41.1 | 780.93 |
| Alzheimer Disease | G30.9, G30.1, G30.0, G30.8 | 331.0, 294.11, 294.10 |
| Parkinsonism | G20, G21.2, G21.19 | 332.0, 294.10, 332.1 |
| Major Depressive Disorder | F32.9, F33.1, F33.2, F41.8, F33.42, F33.41, F33.3, F01.51, F33.9, F43.21, F32.4, Z13.89, F34.1, F31.32, F32.1, F03.90, F32.8, F43.20, F31.30, F31.81, F31.4, F31.9, F32.2, Z63.4, F32.3, F32.5, F32.89, F43.29, F32.0 | 311, 296.32, 296.33, 300.4, 296.36, 296.35, 296.34, 290.43, 296.30, 309.0, 296.25, V79.0, 296.20, 296.52, 296.22, 290.13, 296.50, 296.89, 296.53, 296.80,  290.21, 296.24, 296.21 |
| Obstructive Sleep Apnea | G47.52, G47.33, G47.9, G47.30, G47.8, Z72.820, G47.62, G47.39, G47.31, G47.22, F10.982, G47.37, F51.5, Z76.89, G47.36, F51.8, G47.20, G47.12, G47.34,  G47.21, G47.50, G47.69, G47.10 | 327.42, 327.23, 780.50, 780.57, 307.49, 780.58, V69.4, 327.52, 327.29, 327.32, 327.21, 291.82,  327.27, 307.47, 327.20, 327.30, 780.56, 327.12, 327.24, 327.31, 307.48, 327.26, 780.53 |
