## Supplementary figures and images for "Examining heterogeneity in dementia using data-driven unsupervised clustering of cognitive profiles"

### Supplementary Figure S3

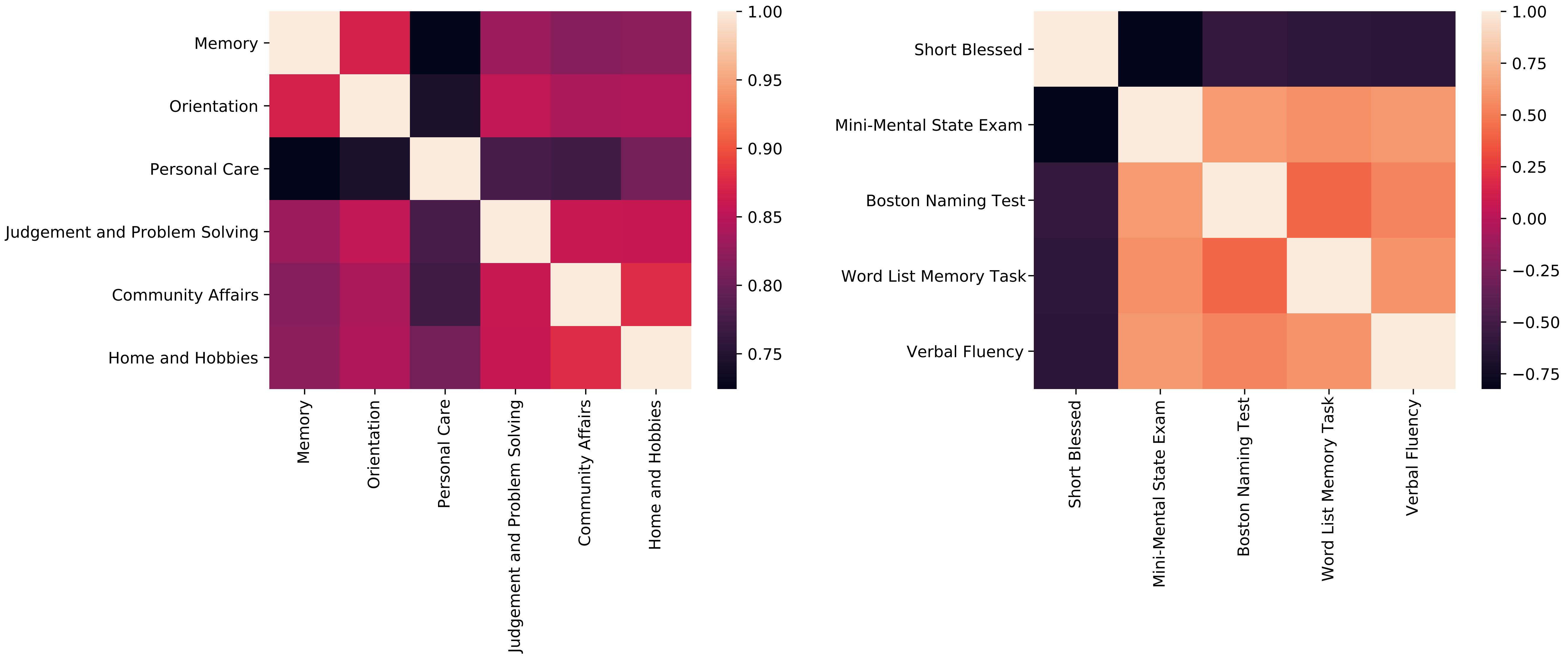
