## Supplementary Table S4 for "Examining heterogeneity in dementia using data-driven unsupervised clustering of cognitive profiles"

|  |  | **Source** | | | | | | | | | | |
| --- | --- | --- | --- | --- | --- | --- | --- | --- | --- | --- | --- | --- |
|  | **Subtype** | **C_2_** | **C_9_** | **C_4_** | **C_7_** | **C_5_** | **C_8_** | **C_6_** | **C_10_** | **C_3_** | **C_1_** | Total |
| **Target** | **C_2_** | **38** | 14 | 3 | 0 | 0 | 0 | 0 | 0 | 1 | 0 | **56** |
|  | **C_9_** | 15 | **75** | 24 | 13 | 7 | 1 | 0 | 1 | 0 | 0 | **136** |
|  | **C_4_** | 2 | 19 | **67** | 56 | 44 | 3 | 5 | 0 | 3 | 0 | **199** |
|  | **C_7_** | 0 | 1 | 7 | **54** | 50 | 15 | 5 | 9 | 8 | 9 | **158** |
|  | **C_5_** | 0 | 2 | 5 | 16 | **75** | 5 | 4 | 8 | 29 | 12 | **156** |
|  | **C_8_** | 0 | 0 | 0 | 8 | 4 | **10** | 2 | 2 | 1 | 0 | **27** |
|  | **C_6_** | 0 | 0 | 0 | 0 | 1 | 3 | **13** | 8 | 4 | 5 | **34** |
|  | **C_10_** | 0 | 0 | 0 | 1 | 2 | 1 | 6 | **12** | 8 | 7 | **37** |
|  | **C_3_** | 0 | 0 | 0 | 0 | 0 | 0 | 1 | 9 | **37** | 19 | **66** |
|  | **C_1_** | 0 | 0 | 0 | 0 | 0 | 0 | 0 | 1 | 3 | **19** | **23** |
